# Fusion protein EWS-FLI1 is incorporated into a protein granule in cells

**DOI:** 10.1101/2020.05.31.122028

**Authors:** Nasiha S. Ahmed, Lucas M. Harrell, Daniel R. Wieland, Michelle A. Lay, Valery F. Thompson, Jacob C. Schwartz

**Affiliations:** Department of Molecular and Cellular Biology, The University of Arizona, Tucson, AZ 85719; Department of Chemistry and Biochemistry, The University of Arizona, Tucson, AZ 85719

**Keywords:** Ewing sarcoma, fusion proteins, phase separation, granules, transcription

## Abstract

Ewing sarcoma is driven by fusion proteins containing a low complexity (LC) domain that is intrinsically disordered and a powerful transcriptional regulator. The most common fusion protein found in Ewing sarcoma, EWS-FLI1, takes its LC domain from the RNA-binding protein EWSR1 (Ewing Sarcoma RNA-binding protein 1) and a DNA-binding domain from the transcription factor FLI1 (Friend Leukemia Virus Integration 1). EWS-FLI1 can bind RNA polymerase II (RNA Pol II) and self-assemble through its low-complexity (LC) domain. The ability of RNA-binding proteins like EWSR1 to self-assemble or phase separate in cells has raised questions about the contribution of this process to EWS-FLI1 activity. We examined EWSR1 and EWS-FLI1 activity in Ewing sarcoma cells by siRNA-mediated knockdown and RNA-seq analysis. More transcripts were affected by the EWSR1 knockdown than expected and these included many EWS-FLI1 regulated genes. We reevaluated physical interactions between EWS-FLI1, EWSR1, and RNA Pol II, and employed a cross-linking based strategy to investigate protein assemblies associated with the proteins. The LC domain of EWS-FLI1 was required for the assemblies observed to form in cells. These results offer new insights into a protein assembly that may enable EWS-FLI1 to bind its wide network of protein partners and contribute to regulation of gene expression in Ewing sarcoma.

## INTRODUCTION

RNA-binding proteins are key players in every step of mRNA biogenesis (Moore and Proudfoot 2009). FUS, EWSR1, and TAF15 comprise the FET family of proteins similar in structure and ubiquitously expressed. These proteins are predominantly nuclear and bind thousands of RNA transcripts with a degenerate specificity (Schwartz et al. 2015; Ozdilek et al. 2017). Although significantly more published work has focused on FUS, recent studies have begun to explore what paradigms and mechanisms of FUS may also apply to EWSR1. FET proteins contribute to the regulation of expression for thousands of genes (Masuda et al. 2015; Schwartz et al. 2012). FUS and EWSR1 bind the C-terminal domain (CTD) of RNA Pol II and modulate its phosphorylation (Kwon et al. 2013; Schwartz et al. 2013; Gorthi et al. 2018). EWSR1 has also been shown to bind a broad network of nascent RNA transcripts, RNA-processing factors, and transcription factors (Chi et al. 2018). The large number of interactions maintained by FET proteins provide a variety of avenues to influence gene expression in cells (Kawaguchi et al. 2020; Chi et al. 2018).

The FET protein genes are frequently involved in genomic translocation events in sarcomas (Riggi et al. 2007; Tan and Manley 2009). The second most common pediatric bone cancer, Ewing sarcoma, is driven by a translocation event fusing the N-terminal low complexity (LC) domain of a FET protein and the DNA-binding domain (DBD) of an ETS transcription factor. In 85% of Ewing sarcomas, a chromosomal translocation produces the EWS-FLI1 oncogene and the resulting fusion protein possessing the LC domain of EWSR1 and the DBD from the FLI1 (Delattre et al. 1992; Grünewald et al. 2018). Ewing sarcomas typically possess few mutations other than the EWS/ETS fusion oncogene, which is sufficient to transform cells (Kovar et al. 1996; Tirode et al. 2014; Stewart et al. 2014). The FLI1 DBD in EWS-FLI1 recognizes a short DNA sequence, GGAA, and binds single sites and microsatellites of minimally 10 repeated motifs (Gangwal et al. 2008; Johnson et al. 2017; Vo et al. 2016). EWS-FLI1 binds RNA Pol II and a number of transcription factors, enhancer, and repressor complexes (Riggi et al. 2014; Boulay et al. 2017; Selvanathan et al. 2019; Theisen et al. 2016; Gorthi et al. 2018). Nevertheless, the mechanism for EWS-FLI1 to change transcription for many hundreds of genes remains poorly understood.

FET proteins have an ability to assemble into higher-order ribonucleoprotein (RNP) assemblies through a process commonly known as phase separation (Kato and McKnight 2018; Springhower et al. 2020; McSwiggen et al. 2019b; Wang et al. 2018; Qamar et al. 2018). RNA seeds this process to trigger FET protein binding to RNA Pol II (Kwon et al. 2013; Schwartz et al. 2013). The LC domain of the FET proteins contains a repeated amino acid motif, [S/G]Y[S/G]. The aromatic sidechain of the tyrosine residues is critical for the domain to undergo phase separation (Kwon et al. 2013; Lin et al. 2017). Phase separation is a process contributing to formation of several RNP granules, such as stress granules, processing (P)–bodies, and nucleoli (Riback et al. 2020). Recent studies offer evidence that phase separation is involved in gene transcription (Wei et al. 2020; McSwiggen et al. 2019a; Guo et al. 2019; Thompson et al. 2018). This mechanism of transcription control likely incorporates FET proteins (Abraham et al. 2020; Chong et al. 2018; Murray et al. 2017).

Proteins fused with the LC domain of EWSR1 form loose interacting assemblies in cells that bind RNA Pol II (Chong et al. 2018; Boulay et al. 2017; Johnson et al. 2017). EWS-FLI1 can form homotypic self-interactions and heterotypic interactions through the LC domain (Spahn et al. 2003). EWS-FLI1 binds and recruits EWSR1 to enhancer regions of the genome (Gorthi et al. 2018; Boulay et al. 2017). Homo-oligomerization through the LC domain is required to stabilize EWS-FLI1 binding at GGAA microsatellites (Johnson et al. 2017). Oligomerization or phase separation appears to be required for EWS-FLI1 to control transcription and initiate transformation, but many questions remain about the form this process takes in cells (Chong et al. 2018; Gorthi et al. 2018; Boulay et al. 2017; Kwon et al. 2013).

Our lab has found the FET protein FUS binds RNA Pol II in a nuclear granule during transcription in cells (Thompson et al. 2018). The importance of oligomerization to FET protein function suggests that EWS-FLI1 may also incorporate into a granule while functioning in cells. Here we investigate EWS-FLI1 associations with protein assemblies or granules in cells. We compared control of gene expression by EWSR1 and EWS-FLI1 and the influence each has on cell transformation. We employed crosslinking-based methods to identify protein assemblies that associate with EWSR1 or EWS-FLI1 in cells and explore their physical characteristics. Our aim was characterizing assemblies incorporating the fusion protein to reveal greater understanding of the complex mechanisms used by EWS-FLI1 to direct cell transcription and transformation.

## RESULTS

### Transcript levels in Ewing sarcoma are affected by both EWSR1 and EWS-FLI1

EWSR1 and EWS-FLI1 share the same LC domain, which undergoes oligomerization for regulating transcription (Kwon et al. 2013; Chong et al. 2018). These studies show homotypic self-interactions by the LC domain of EWS-FLI1 are required to its activity toward transcription and transformation. On the other hand, EWS-FLI1 has a dominant negative effect on some EWSR1 activities in Ewing sarcoma (Gorthi et al. 2018; Embree et al. 2009). Since EWS-FLI1 also recruits EWSR1 to GGAA-rich sites along chromosomal DNA, we performed RNA-seq to investigate the effects of EWSR1 on genes repressed or activated by EWS-FLI1.

Poly-adenylated RNA was isolated from the Ewing sarcoma cell line, A673, which expresses EWS-FLI1 and EWSR1. The mRNA for EWS-FLI1, EWSR1, or both was knocked down using small interfering RNAs (siRNA). An siRNA specific for EWS-FLI1 (siEF) targeted the 3’ region of the EWS-FLI1 mRNA that is not found in the EWSR1 transcript. The siRNA specific for EWSR1 (siEWSR1) targeted to the 3’ region of its mRNA. Both transcripts were knocked down by an siRNA (siE-EF) targeting the 5’ region of both transcripts (**Supplemental Figure S1A**). The specificity and protein reduction produced by the siRNA knockdowns were confirmed by western analysis (**Supplemental Figure S1B**).

Analysis of expressed A673 transcripts (n = 10706) by RNA-seq revealed large numbers that were significantly increased (n = 1537) or decreased (n = 1282) by >1.6-fold (p-adj < 0.05). Compared with two previously published studies in A673 or SK-N-MC cells, we found 911 transcripts affected by the EWS-FLI1 knockdown in our study were also affected in all three published datasets (Riggi et al. 2014; Sankar et al. 2013). We also inspected a signature set of 148 genes previously identified as repressed targets of EWS-FLI1 in Ewing sarcoma (Smith et al. 2006). Of the 147 identified in our transcriptome, the knockdown by siEF significantly increased transcript levels for 133 (>1.6-fold, p-adj < 0.05, **Figure 1A, Supplemental Table 2**).

**Figure 1.**
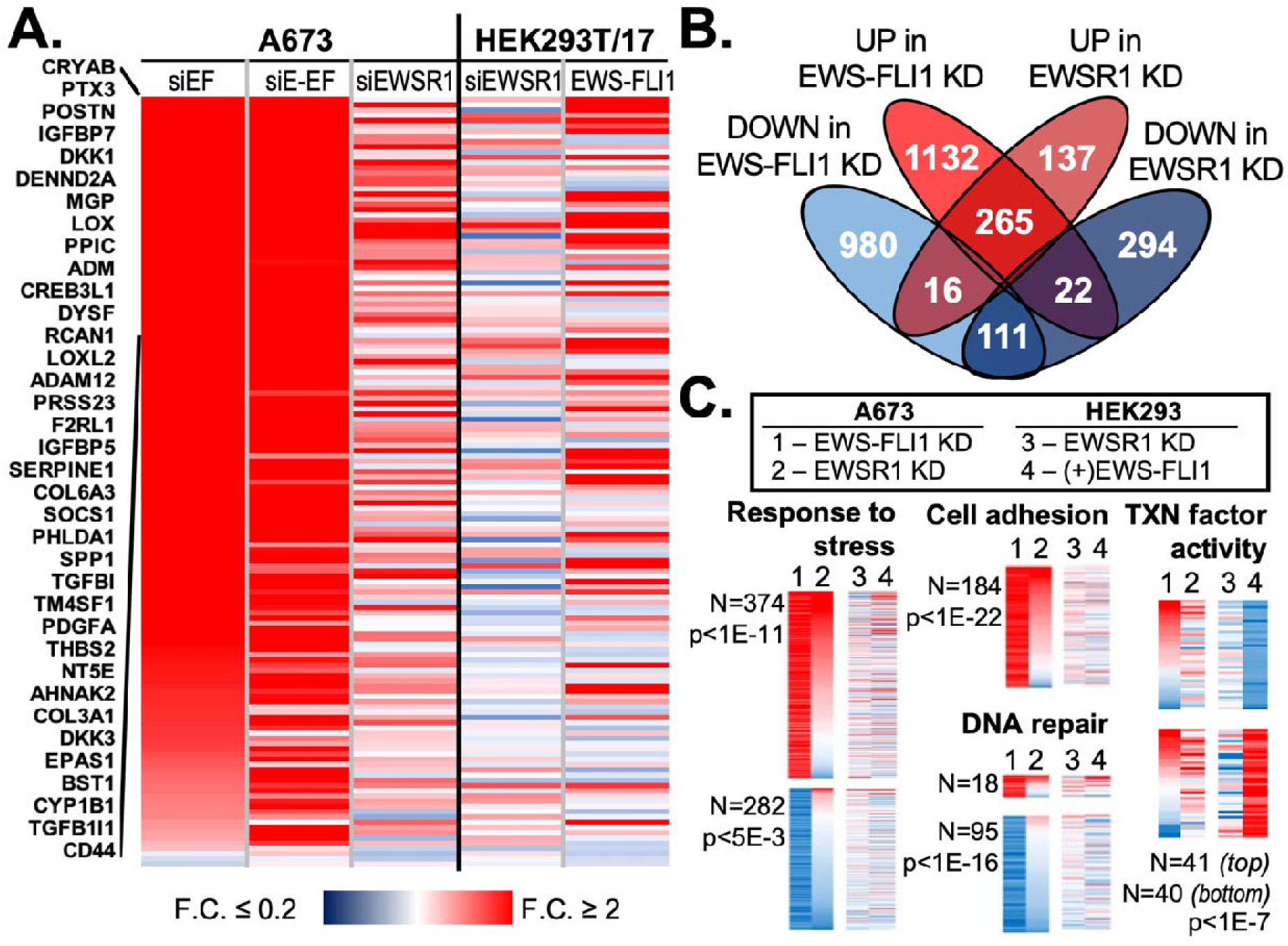
Many transcripts are similarly affected by knockdown of EWS-FLI1 or EWSR1 in Ewing sarcoma. (**A**) A heat map shows fold changes for the 147 signature genes repressed by EWS-FLI1 in A673 cells. Results are sorted according to change in the siEF treatment, with names of the 39 most affected gene shown. Changes to mRNA levels in HEK293T/17 after knockdown by siEWSR1 or expression of EWS-FLI1 in HEK293T/17 cells are also shown. (**B**) The Venn diagram shows overlaps in transcripts increased or decreased by the knockdown (KD) of EWS-FLI1 or EWSR1 in A673 cells. The threshold required for increased or decreased transcripts was 1.6-fold change and adjusted p-value, p-adj < 0.05. (**C**) Heat maps show changes in transcript abundances in A673 after EWS-FLI1 (1) or EWSR1 (2) knockdown or in HEK293T/17 after EWSR1 knockdown (3) or EWS-FLI1 (4) expression. Included genes were taken from selected GO associations identified among those affected by the EWS-FLI1 knockdown: response to stress (GO:0006950), cell adhesion (GO:0007155), and DNA repair (GO:0006281). These are sorted by changes from EWS-FLI1 (1) then EWSR1 (2) knockdown. Changes of genes associated with transcription factor activity (GO:0003700) are shown for those affected by exogenous EWS-FLI1 in HEK293T/17 cells. The number of genes plotted is indicated and adjusted p-values corrected by the Benjamini & Hochberg False Discovery Rate.

Following the knockdown of EWSR1, relatively fewer gene transcripts increased (n = 418) or decreased (n = 427) in abundance. Consistent with this, the distance of siEF from siSCR samples in a principal component analysis was much greater than that for the siEWSR1 treated cells (**Supplemental Figure S1C–D**). Of the 147 repressed genes, 43 increased in transcripts abundance after EWSR1 knockdown (>1.6-fold change, p-adj < 0.05, **Figure 1A**). Most transcripts increased by EWSR1 knockdown also increased after EWS-FLI1 knockdown (n = 265, **Figure 1B**). Only 25% (n = 111) of gene transcripts reduced by knockdown of siEWSR1 were also reduced by EWS-FLI1 knockdown. Very few genes diverged in response to the two knockdowns (n = 22 decreased and 16 increased by siEWSR1, **Figure 1B**). We also analyzed RNA-seq results following the knocked down by siE-EF, finding 859 increased and 635 decreased transcripts in common with knockdown by siEF. Most genes significantly changed by siEF and siEWSR1 were also increased (n = 190) or decreased (n = 84) by knockdown with siE-EF (**Supplemental Table 2**).

A previous study of an EWSR1 knockdown by a stable shRNA method caused few effects on transcript levels in A673 cells (Sankar et al. 2013). We included in our analysis the data made publicly available by the published study, which also identified only 129 expressed transcripts affected by the EWSR1 knockdown. Of these, 32 transcripts were affected by the EWSR1 knockdown in our experiment (**Supplemental Table 2**). The reduction in EWSR1 transcript did not differ between the shRNA or siRNA treated samples. However, we noted the shRNA treated A673 cells were cultured for 2 weeks under selection by antibiotics (Sankar et al. 2013). We hypothesized this time in culture may allow enrichment in the cell population of those with restored EWS-FLI1 activity and regulation of gene expression by a mechanism circumventing the role contributed by EWSR1.

We considered whether effects of the EWSR1 knockdown would differ in the absence of EWS-FLI1. We chose a non-Ewing cell line, HEK293T/17, to perform an RNA-seq analysis after knockdown of EWSR1 or exogenous expression of EWS-FLI1 from a transfected plasmid. The EWSR1 knockdown observed by western analysis was comparable to that of A673 cells (**Supplemental Figure S1E**). We found 98% of 10706 genes analyzed for A673 cells above met the same requirement to be categorized as expressed in HEK293T/17 cells.

This was also found true for 97% of genes with transcripts affected by the EWS-FLI1 knockdown in A673. Compared to results for Ewing sarcoma cells, the EWSR1 knockdown in HEK293T/17 cells caused levels of relatively few transcripts to significantly increase (n = 139) or decrease (n = 148) by >1.6-fold (**Supplemental Table 2, Supplemental Figures S1F–G**). Of 95 genes affected in both cell lines, two-thirds (n = 62) decreased in transcript abundance (**Supplemental Figure S1H**). Additionally, 23 of the 33 genes whose transcripts increased in both cell lines after EWSR1 knockdown, were also increased in transcript levels following EWS-FLI1 knockdown in A673.

Compared to knocking down endogenous EWS-FLI1, exogenous EWS-FLI1 expression in HEK293T/17 cells caused fewer mRNA transcripts to be significantly increased (n = 509) or decreased (n = 389) in abundance by >1.6-fold and p-adj < 0.05 (**Supplemental Figure S1F–G**). Many signature genes (n = 29) repressed in Ewing sarcoma were not silenced, but exogenous expression of EWS-FLI1 resulted in increases to their transcript abundances (**Figure 1A**). Of the transcripts affected in both cell lines, most of those increased by exogenous EWS-FLI1 were repressed by EWS-FLI1 in A673 cells (n = 163 or 75% of 219 genes affected, **Supplemental Figure S1H**). We considered whether divergent effects of EWS-FLI1 may relate to differences in expression levels for each cell line and found the average transcript levels in A673 were 5-fold higher than in HEK293T/17 cells for genes activated by the knockdown or exogenous expression of EWS-FLI1 (p < 0.05, Student’s t-test with equal variances). This observation suggested that exogenous EWS-FLI1 may bring HEK293T/17 transcript levels closer to those in A673. Most genes silenced by exogenous EWS-FLI1 agreed with knockdown results (n = 82 of 125 affected genes, **Supplementary Figure S1H**).

Finally, we investigated the gene ontology (GO) associations among genes affected by EWS-FLI1 or EWSR1. Those found among genes increased by EWS-FLI1 knockdown included cell adhesion (n = 184, p-adj < 1E-22), response to stress (n = 374, p-adj < 1E-11), and cell death (n = 102, p-adj < 2E-2) (**Figure 1C, Supplemental Figure S2**). After EWSR1 knockdown, the number of adhesion genes whose transcripts increased was similar in proportion to increases caused by the EWS-FLI1 knockdown (68% of 113 affected) and higher for genes associated with stress (65% of 272 affected) and cell death (69% of 84 affected). Examples of GO associations among genes silenced by the EWS-FLI1 knockdown included DNA repair (n = 95, p-adj ≤ 1E-16), DNA damage response (n = 125, p-adj ≤ 1E-19), and cell cycle (n = 180, p-adj ≤ 1E-42) (**Figure 1C, Supplemental Figure S2**). Exogenous EWS-FLI1 mostly increased DNA damage response gene transcripts (84% of 32 affected) but affected few cell cycle genes (n = 15). The EWSR1 knockdown effects on DNA repair genes were mixed (n = 51), but mRNA levels were mostly reduced for affected cell cycle genes (72% of 43 affected). Knockdown also suggested that EWS-FLI1 and EWSR1 activity increased expression of genes involved in chromatin modifications, including histone acetyltransferases and methyltransferases (**Supplemental Figure S2**).

Regulators of gene expression were affected by both EWS-FLI1 knockdown (n = 380) and exogenous expression (n = 111). An example of agreement between these treatments was noted for the 41 transcription factor genes silenced by EWS-FLI1 in HEK293T/17, while agreement was less for the 40 transcripts whose levels increased (**Figure 1C, Supplemental Figure S2**). The knockdown of EWS-FLI1 affected binding to unfolded proteins (n = 38, p ≤ 2E-3). Lastly, we noted effects on cellular organelle genes. Knockdown by siEF or siE-EF reduced expression for 21 of 46 genes associated with the nuclear pore and 204 nucleolar genes (**Supplemental Figure S2**).

In summary, the EWSR1 knockdown in Ewing sarcoma yielded similar effects as knocking down EWS-FLI1 and different from an EWSR1 knockdown in a non-Ewing cell line. This supports the hypothesis that a functional interaction between the two proteins may be present in Ewing sarcoma (Gorthi et al. 2018; Li et al. 2007). However, the diminished effects of a loss of EWSR1 found by the lengthier shRNA knockdown protocol motivated us to pursue additional approaches to test potential functions shared by EWSR1 and EWS-FLI1 in Ewing sarcoma.

### EWSR1 is required for anchorage-independent growth in Ewing sarcoma

The loss of EWS-FLI1 can block anchorage-independent cell growth for Ewing sarcoma cell lines in soft agar colony formation assays (Smith et al. 2006; Chaturvedi et al. 2012). We hypothesized that an interaction between EWS-FLI1 and EWSR1 that was functional and significant to Ewing sarcoma biology may be reflected by changes to the tumorigenic capacity of cells in response to the knockdown of either protein.

We performed soft agar assays for A673 cells after EWSR1 or EWS-FLI1 knockdown. The knockdown of each protein was confirmed to be efficient and specific by western blot analysis (**Figure 2A**). We monitored cell growth and observed no change in cell count over a period of 4 days after each knockdown or the control siSCR treatment (**Supplemental Figure S3A**). Transfected A673 cells seeded on soft agar were grown for 3 to 4 weeks. The knockdown of EWS-FLI1 resulted in 60% fewer colonies relative to siSCR controls (n = 3, *p* = 0.04, Student’s t-test, **Figure 2B**). Cells with the EWSR1 knockdown yielded 90% fewer colonies, revealing a dependency on EWSR1 for growth on soft agar (n = 3, *p* = 0.004, Student’s *t*-test).

**Figure 2.**
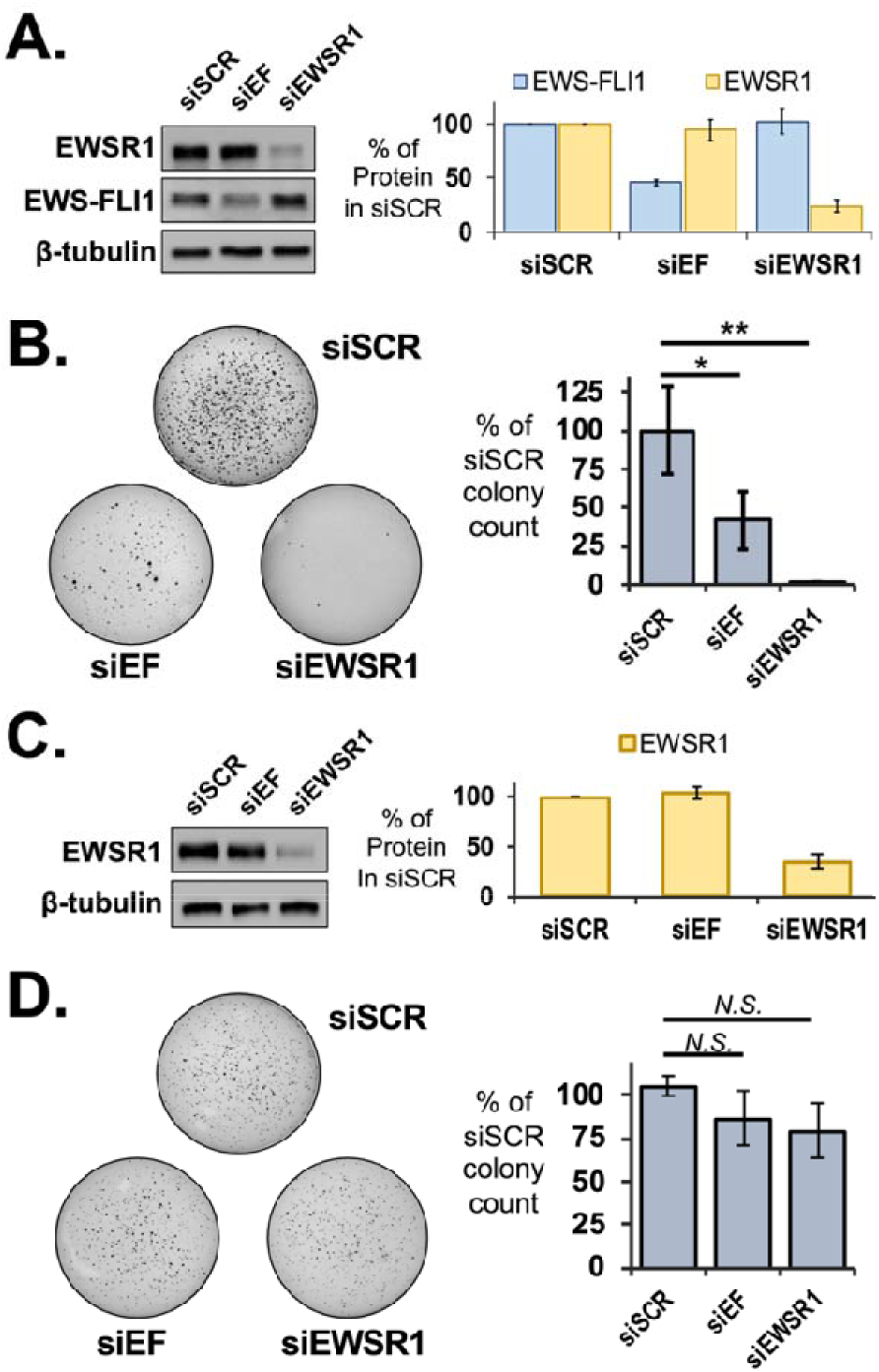
Loss of EWSR1 inhibits anchorage-independent growth in Ewing sarcoma cells. (**A**) Left, western assays reveal knockdown for EWSR1 or EWS-FLI1 in A673 cells for treatment with siSCR, siEF, or siEWSR1. Right, averaged levels of protein were determined by densitometry relative to siSCR treatments (n = 3). (**B**) Left, soft agar assays were performed with A673 cells treated with siSCR, siEF, or siEWSR1. Right, colonies were counted to reveal reductions in averaged values relative to siSCR treatment (n = 3). (**C**) Left, western assays reveal knockdown for EWSR1 or EWS-FLI1 in HEK293T/17 cells for treatment with siSCR, siEF, or siEWSR1. Right, averaged levels of protein were determined by densitometry relative to siSCR treatments (n = 4). (**D**) Left, soft agar assays were performed for HEK293T/17 cells treated with siSCR, siEF, or siEWSR1. Right, colonies were counted to reveal no change in averaged values relative to siSCR treatment (n = 4). All error bars represent standard deviation. Student’s *t*-test was calculated assuming equal variances: ** *p* < 0.01; * *p* < 0.05; n.s., not significant (*p* > 0.05).

These results suggested that the loss of EWSR1 or EWS-FLI1 led to a similar loss in tumorigenic capacity for Ewing sarcoma cells. We repeated the soft agar assays using SKNMC cells. By western blot analysis, the reduction of EWS-FLI1 or EWSR1 protein was found to be similar for knockdowns in SKNMC and A673 cells (**Supplemental Figure S3B**). In SKNMC cells, the EWS-FLI1 knockdown resulted in 80% fewer colonies and 90% fewer colonies for the EWSR1 knockdown (**Supplemental Figure S3C**). The EWSR1 knockdown might not be expected to affect tumorigenic capacity since the shRNA knockdown of EWSR1 found few effects on EWS-FLI1 target genes (Sankar et al. 2013). Conversely, the effects that siEWSR1 produced for EWS-FLI1 target genes would raise the expectation that either knockdown can yield the same reduction in cell growth we have observed in soft agar assays.

The loss in colony growth after an EWSR1 knockdown in A673 cells could be interpreted as unrelated to EWS-FLI1 if a cell line without the fusion protein saw the same result. We chose a non-Ewing cell line, HEK293T/17 cells, to test the effects for the EWSR1 knockdown in a background without EWS-FLI1 or detectable FLI1 protein. Transfection with siEWSR1 effectively diminished EWSR1 protein but transfection with siEF had no effect (**Figure 2C**). HEK293T/17 cells treated with siEWSR1, siEF, or the siSCR control produced no difference in colony numbers on soft agar (n = 4, *p* > 0.05, Student’s *t*-test, **Figure 2D**). This result left the possibility that EWS-FLI1 may impose a new role on EWSR1 in controlling cell phenotypes related to tumorgenicity.

Observing that an EWSR1 knockdown did not affect HEK293T/17 growth on soft agar, we asked whether the fusion protein could recreate the sensitivity to the loss of EWSR1 observed in Ewing sarcoma cells. We expressed V5-tagged EWS-FLI1 protein in HEK293T/17 cells (**Supplemental Figure S4A**). The expression of the fusion protein from a transfected plasmid did not inhibit growth of HEK293T/17 in the soft agar assay compared to an empty plasmid (**Figure 3A**). However, the knockdown of EWSR1 in HEK293T/17 cells expressing the fusion protein yielded 60% fewer colonies compared with the siSCR-treated control (n = 6, *p* < 0.001, Student’s *t*-test, **Figure 3B**). The LC-domain shared by EWS-FLI1 and EWSR1 mediates homotypic interactions and binding to each other in cells (Chong et al. 2018; Gorthi et al. 2018; Boulay et al. 2017; Spahn et al. 2002). We confirmed this for V5-EWS-FLI1 expressed in these HEK293T/17 cells using a co-immunoprecipitation (co-IP) assay. With an EWSR1 specific antibody (B-1), we found by western assay V5-EWS-FLI1 protein eluted with EWSR1 and not in the control samples for an IP with non-specific IgG (**Supplemental Figure S4B**). As noted by previous studies, we found co-IP experiments in A673 were inconsistent in recovering endogenous EWS-FLI1 enriched by an EWSR1 IP above controls (**Supplemental Figure S4C**) (Spahn et al. 2003). We likewise interpreted the result to be influenced by a lower abundance of endogenous EWS-FLI1, the weakness in binding for homotypic and heterotypic interactions, or both.

**Figure 3.**
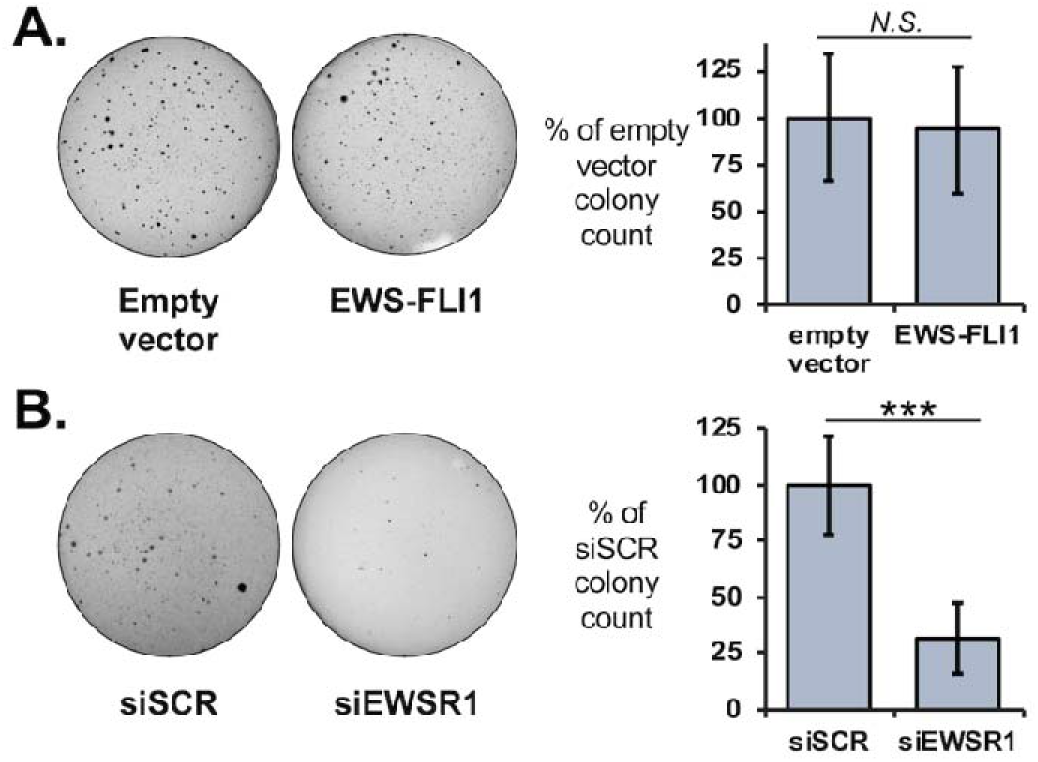
EWS-FLI1 can alter effects on cell growth for an EWSR1 knockdown. (**A**) Left, growth on soft agar was assessed for HEK293T/17 cells after transfection of empty plasmids or a plasmid expressing V5-EWS-FLI1. Right, colonies were counted to reveal no change in averaged values relative to siSCR treatment (n = 4). (**B**) Left, soft agar assays were performed with HEK293T/17 cells expressing V5-EWS-FLI1 and co-transfected with siSCR or siEWSR1 (n = 6). Right, colonies were counted to reveal a reduction in averaged values relative to siSCR treatment. Error bars represent standard deviation. Student’s *t*-test was calculated for assuming equal variances: \*\*\**p* < 0.001.

### EWSR1 and RNA Pol II associate with protein granules in cells

The LC domain shared by FET proteins and fusion proteins driving Ewing sarcoma are capable of undergoing phase separation, a process that can form granules in cells (Kato et al. 2012; Kato and McKnight 2018; Lin et al. 2015). Consistent with the weak protein interactions for *in vitro* phase separation, granules in cells do not maintain their integrity after cell lysis (Wang et al. 2018; Qamar et al. 2018). Homotypic interactions for FET proteins have been confirmed in cells by previous studies (Spahn et al. 2003; Boulay et al. 2017; Kwon et al. 2013). To date, studies have not definitively established one of the several assemblies observed *in vitro* to be the predominant form in cells. These include (1) large multimer assemblies but of ordinary size for protein complexes in cells, (2) simple polymer or fiber-like assemblies, (3) large assemblies formed of relatively stable interactions described as hydrogels, or (4) condensates formed of weak interactions that give rise to liquid-like properties. We have previously developed protocols that stabilize weak protein interactions for FET proteins with formaldehyde crosslinking (Thompson et al. 2018). With this approach, we characterized a granule formed by RNA Pol II transcription in cells. We now addressed whether EWSR1 associates with large macromolecular assemblies or condensates in cells.

We applied our SEC-based approach using lysates prepared from HEK293T/17 cells to separate large protein assemblies from ordinary molecular complexes and quantify amounts of EWSR1 and RNA Pol II in each fraction (Thompson et al. 2018). Important steps to the interpretation of our analysis by crosslinking include that all samples were subjected to sonication and nuclease treatment to minimize detection of assemblies tethered by nucleic acids rather than direct protein-protein interactions. Analysis using dynamic light scattering (DLS) indicated that SEC with a CL-2B column could separate particles from 14 nm to 150 nm in diameter (**Figure 4A**) (Thompson et al. 2018). The expected size of RNA Pol II is 25 nm at its longest axis. Particles of this size were observed at a 20 mL elution volume (**Figure 4A**, grey dashed line). UV absorption signals were maximal at >20 mL elution volumes, indicating most protein complexes or monomers were < 25 nm in diameter (n = 3, **Supplemental Figure S5A**). Lysates of crosslinked cells were prepared using buffers containing 6 M urea to prevent non-specific interactions not crosslinked in the cell. The UV trace for crosslinked samples did not differ appreciably from those with no crosslinking, indicating most proteins were not crosslinked in large assemblies (n = 3, **Supplemental Figure S5B**).

**Figure 4.**
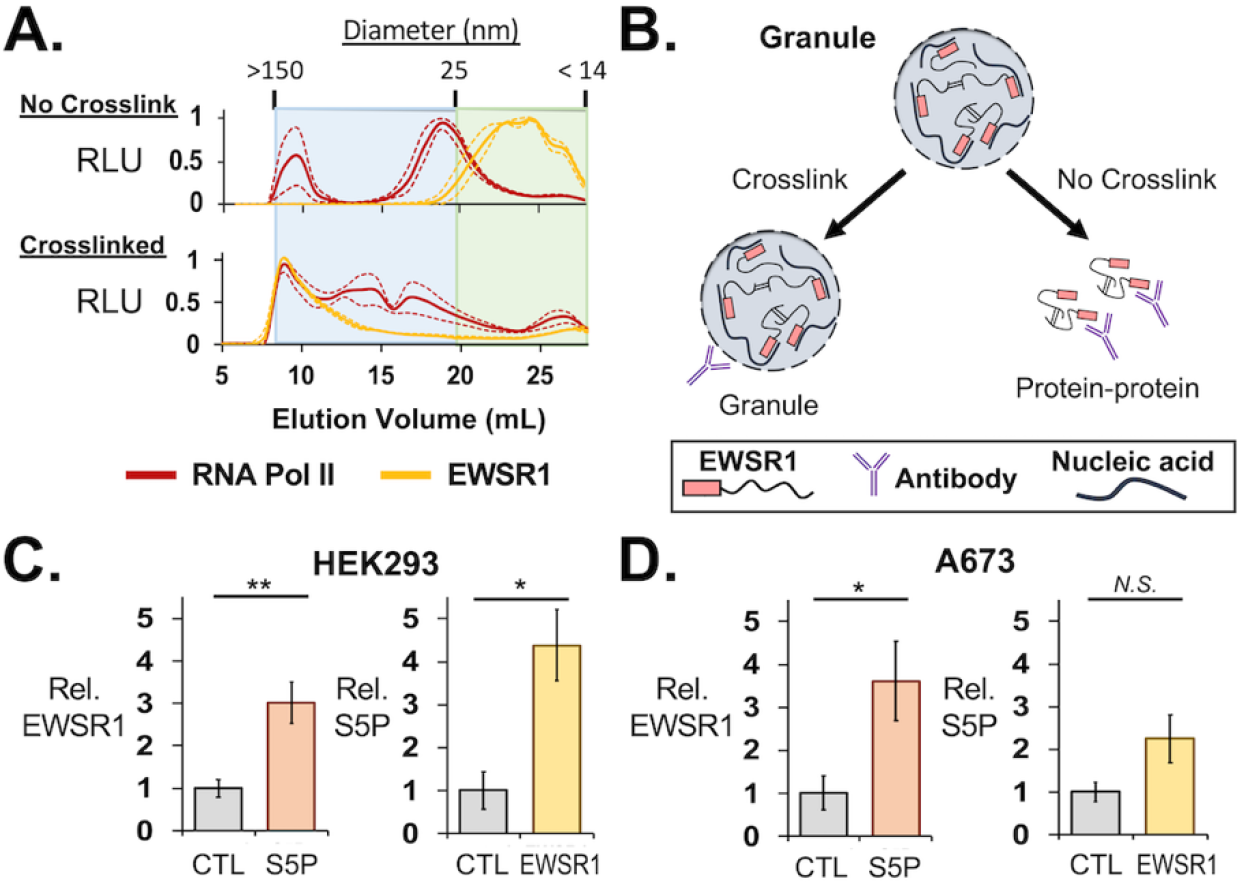
EWSR1 and RNA Pol II are assembled in large protein assemblies. (**A**) Total protein from HEK293T/17 cell lysates was analyzed by SEC to separate proteins, complexes, or assemblies by size. The averaged ELISA signals are shown normalized to their maximum values for EWSR1 (C-9 antibody), yellow, or RNA Pol II (CTD4H8 antibody), red, as measured by ELISA. For samples from cells not crosslinked, top, RNA Pol II signals were highest in fractions eluting just before 20 mL, corresponding to particles of 25 nm in size. EWSR1 eluted after 20 mL or as a particle < 25 nm in size. For crosslinked samples, bottom, the elution of EWSR1 and RNA Pol II shifted to early fractions with particles measured to be up to 150 nm in diameter. Dashed lines represent standard error about the mean (n = 3). (**B**) IP assays using cells without crosslinking enrich for ordinary and stable molecular complexes. Those using crosslinked cells can recover large molecular assemblies that form by weak protein interactions. (**C**) Left, ELISA assays detected EWSR1 eluted from crosslinked IP assays of phosphorylated (S5P) RNA Pol II (Abcam, ab5131) in HEK293T/17 cells (n = 3). Right, S5P RNA Pol II was detected for crosslinked IP of EWSR1 (B-1 antibody) (n = 3). (**D**) Left, ELISA assays in A673 cells also detected EWSR1 eluted with S5P RNA Pol II. Right, S5P RNA Pol II was detected in crosslinked IP assays of EWSR1 (n = 3). All error bars represent standard error about the mean. AU = absorbance units. RLU = relative luminescence units. Student’s *t*-test was calculated assuming equal variances: ** *p* < 0.01; * *p* < 0.05; *n.s*., not significant (*p* > 0.05).

We performed ELISA assays on the SEC fractions collected using antibodies for RNA Pol II (CTD4H8) and EWSR1 (C-9). As we have previously reported, RNA Pol II eluted just before and near 20 mL, suggesting a particle somewhat larger than 25 nm in diameter (**Figure 4A**, top). The sum of signals quantitated by ELISA indicated that 25% of RNA Pol II was present in particles >50 nm in diameter for cells not treated with formaldehyde. SEC of lysates prepared from crosslinked cells indicated 54% of the total RNA Pol II detected eluted at volumes < 15 mL, for which particle were >50 nm in diameter (**Figure 4A**). EWSR1 was observed to elute in volumes >20 mL, indicating either small complexes or monomers were present in lysates not crosslinked. In crosslinked cells, 71% of the total EWSR1 detected was found in early fractions for particles of at least 50 nm in diameter. We have previously determined by DLS measurements, the first SEC fraction yielded a signal of 150 nm as the diameter of particles at this void volume of the column. Lysates were passed through a 0.45 μm filter before SEC and by TEM, the void fraction was observed to contain particles of up to 300 nm (Thompson et al. 2018). This suggests that particles eluting early may be bigger than 150 nm but are unlikely to be much bigger than 0.3 μm.

We tested whether interactions between EWSR1 and RNA Pol II were crosslinked in HEK293T/17 cells. This approach allows robust detection for weak interactions that would be rarely observed in standard IP protocols, and can be treated with strong detergents to prevent detection of non-specific interactions by these aggregation prone disordered proteins (**Figure 4B**). For IP experiments in HEK293T/17 cells, an antibody specific for the serine-5 phosphorylated (S5P) form of RNA Pol II (Abcam, ab5131) and was found to yield more efficient protein in the IP elution. An antibody specific for the EWSR1 C-terminus (B-1) was used for IP experiments. Significant enrichment for EWSR1 eluting with the polymerase was observed in ELISA assays, relative to a negative control using protein G beads (n = 3, *p* = 0.009, Student’s *t*-test, **Figure 4C**, left). An IP for EWSR1 similarly found enrichment for S5P RNA Pol II (n = 3, *p* = 0.02, Student’s *t*-test, **Figure 4C**, right). We repeated the IP assays using crosslinked A673 cells, which confirmed crosslinking of interaction of S5P RNA Pol II with an EWSR1 IP (n = 3, *p* = 0.04, Student’s *t*-test, **Figure 4D**). The interaction with EWSR1 in A673 cells was enriched in an S5P RNA Pol II IP but without reaching significance (n = 3,*p* > 0.05, Student’s *t*-test).

### EWS-FLI1 is found in a protein granule in cells

We lastly investigated the interactions and cellular assemblies occupied by EWS-FLI1. When observed *in vitro*, higher order assemblies for FET and other LC domain proteins have been observed in a number of alternative structures that vary in appearance and physical characteristics (Qamar et al. 2018; Lin et al. 2015; Kwon et al. 2013; Schwartz et al. 2013). Under electron microscopy, ordinary protein complexes are visibly distinct from amyloid-like fibers, amorphous aggregates, or the rounded structures of the phase separation process that can form a granule in cells. We now sought to characterize EWS-FLI1 associations with RNA Pol II protein assemblies and observe whether EWS-FLI1 or RNA Pol II assemblies contrasted in their physical characteristics that might indicate similar or wholly independent assembly processes.

We tested whether EWS-FLI1 was crosslinked with assemblies of EWSR1 and RNA Pol II. Analogous to the approach used for analysis by SEC, crosslinked cells were lysed in the presence of 1% SDS to ensure interactions detected were due to crosslinks made in the cell rather than non-specific or aggregation after lysis. Steps were included to cleave interactions tethered by nucleic acids to enrich for protein-protein crosslinked interactions. HEK293T/17 cells were transfected with V5-tagged EWS-FLI1 for use in co-IP assays. Antibodies binding the C-terminal portions of either EWSR1 (B-1) or EWS-FLI1 (Abcam, ab15289) allowed specificity for the assay in A673 cells. For RNA Pol II, the CTD4H8 antibody from mouse was used to avoid interference with EWS-FLI1 antibody signals. By ELISA, EWS-FLI1 was found highly enriched in EWSR1 IP samples compared with negative protein G bead controls (n = 3, *p* = 0.01, Student’s *t*-test, **Figure 5A**). EWS-FLI1 was likewise enriched in RNA Pol II IP experiments compared with negative controls (n = 3, *p* = 0.03, Student’s *t*-test, **Figure 5A**). EWSR1 (C-9) and RNA Pol II (CTD4H8) was also found in the IP of EWS-FLI1 expressed in HEK293T/17 cells (**Supplemental Figure S6A** and **S5B**).

**Figure 5.**
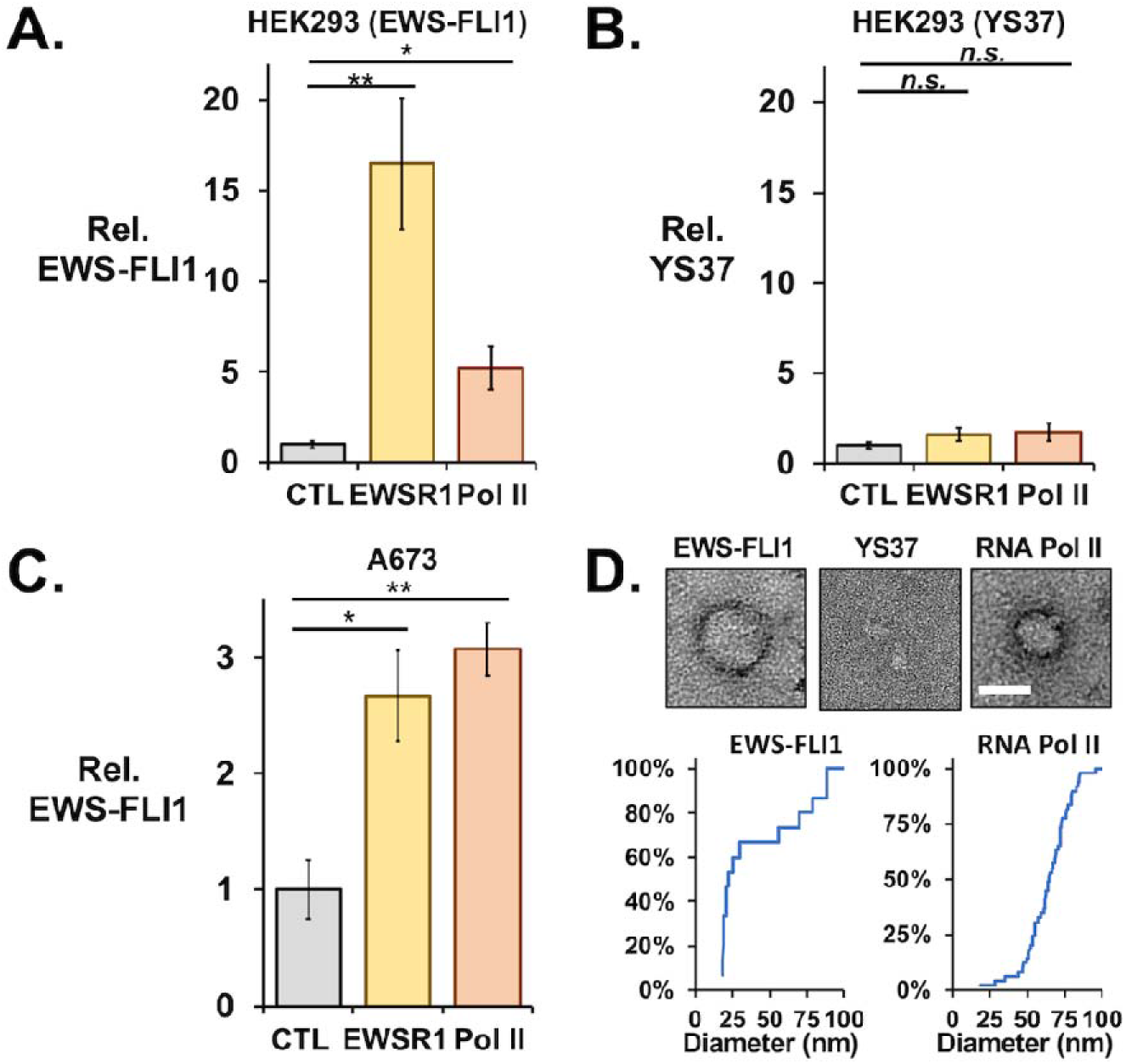
EWS-FLI1 and RNA Pol II coimmunoprecipitate in crosslinked protein granules. ELISA assays using an antibody to FLI1 measured interactions with FLI1-fusion proteins recovered by co-IP assays from crosslinked cells. The IP of EWSR1 (B-1) or RNA Pol II (CTD4H8) recovered V5-EWS-FLI1 (**A**) but not YS37 (**B**) protein expressed in HEK293T/17 cells by plasmid transfections (n = 3 each). (**C**) Crosslinked IP assays of EWSR1 and RNA Pol II also recovered endogenous EWS-FLI1 from A673 cells (n = 4). (**D**) Transmission electron microscopy detected protein particles recovered from crosslinked HEK293T/17 cells by IP of EWS-FLI1 (anti-FLI1 antibody, left) and RNA Pol II (CTD4H8, left), but not of YS37 (anti-FLI1 antibody, center). Scale bar inset represents 50 nm. Cumulative plots show diameters for particles imaged in IP samples for EWS-FLI1 (n = 24) or RNA Pol II (n = 77). All error bars represent standard error about the mean. Student’s *t*-test was calculated assuming equal variances: ** *p* < 0.01; * *p* < 0.05; n.s., not significant (*p* > 0.05).

We confirmed that interactions stabilized by crosslinking were dependent on the key tyrosine residues of the LC domain by expressing an V5-tagged EWS-FLI1 protein with 37 tyrosine residues in the [S/G]Y[S/G] motif replaced by serine (YS37, *provided by the lab of M. Rivera*) (Boulay et al. 2017). We transfected this construct in HEK293T/17 cells and found expression comparable to that of wild-type V5-EWS-FLI1 (**Supplemental Figure S6C** and **S5D**). For crosslinked HEK293T/17 cells, EWSR1 or RNA Pol II did not pulldown the YS37 fusion protein (n = 3, *p* > 0.05, Student’s *t*-test, **Figure 5B**). The reciprocal IP of YS37 also did not find enrichment of EWSR1 or RNA Pol II (**Supplemental Figure S6E** and **S5F**). This result suggested that stabilizing protein interactions by crosslinking could not provide evidence of an EWSR1 or RNA Pol II interaction with an EWS-FLI1 unable to homo-oligomerize through the LC domain. In contrast to our result without crosslinking, we could observe EWSR1 and RNA Pol II in the IP of endogenous EWS-FLI1 from crosslinked A673 cells. Similarly, EWS-FLI1 eluted with EWSR1 or RNA Pol II at significant levels over negative controls when stabilized by crosslinking (n = 4, *p* < 0.05, Student’s *t*-test, **Figure 5C**).

We have previously reported imaging by transmission electron microscopy (TEM) of crosslinked particles associating with RNA Pol II in cells (Thompson et al. 2018). These particles typically appeared round and 50 nm in diameter but could measure up to 270 nm. Their circular shapes could indicate a sphere when in solution, as expected for particles of weak interactions, including liquid-like condensates. We investigated whether the crosslinked assemblies in HEK293T/17 cells shared these properties. By negative-stained TEM, an RNA Pol II IP from crosslinked HEK293T/17 cells yielded round particles up to 95 nm in diameter. Particles sizes averaged 64 ± 15 nm in diameter (n = 77, **Figure 5D**). We imaged samples treated with proteinase K, which eliminated the round particles observed and indicated they were comprised of protein (**Supplemental Figure S7**).

For crosslinked HEK293T/17 cells that expressed V5-EWS-FLI1, we imaged by TEM particles that were also round in shape and varied in size, with an average diameter of 42 ± 27 nm (n = 24, **Figure 5D**, left). To test whether the role of the LC domain, we repeated the assay and TEM imaging using crosslinked HEK293T/17 cells that expressed the YS37 variant of EWS-FLI1. The YS37 IP did not recover round particles visible to TEM (**Figure 5D**). We concluded that protein assemblies associated with EWS-FLI1 were similar shape and comparable size to those observed for RNA Pol II. Their assembly depended on the oligomerization domain of EWS-FLI1. These physical properties observed of particles associated with EWS-FLI1 or RNA Pol II in cells appear consistent with condensates and did not indicate substantial differences in their structural makeup or suggest they had formed by unrelated molecular processes.

## DISCUSSION

The N-terminal domains of FET proteins allow them to oligomerize with disordered domains of similar low complexity and amino acid composition (Murray et al. 2017; Schwartz et al. 2015). We compared the roles of EWSR1 and EWS-FLI1 to control gene expression and found that EWS-FLI1 and EWSR1 can regulate transcription for hundreds of identical gene targets. EWSR1 and the fusion protein each affect the signature property of transformation, anchorage-independent cell growth. The same deficiency in growth on soft agar indicates a functional interaction but without distinguishing whether this reflects protein-protein binding. The same assay for a non-Ewing cell line is not affected by the EWSR1 knockdown, but the effect was reproduced when the fusion protein was introduced. This result indicates that interactions between EWS-FLI1 and EWSR1 are significant enough to alter the cellular function of these proteins.

The mechanisms by which FET proteins and their fusion products interact with RNA Pol II have the potential to influence paradigms for transcription regulation in higher eukaryotes (Abraham et al. 2020; Zamudio et al. 2019; Chong et al. 2018; Thompson et al. 2018). FET proteins bind the polymerase *in vitro* while oligomerized into large assemblies (Schwartz et al. 2013; Kwon et al. 2013). Both FUS and EWSR1 can alter phosphorylation of RNA Pol II at positions Ser2 or Ser5, respectively, in the repeated heptad motif of the CTD (Schwartz et al. 2012; Gorthi et al. 2018). Phase separation associated with EWSR1 activity and RNA Pol II transcription invites the question whether this is a feature of EWS-FLI1 in cells and if so, how comparable is an EWS-FLI1 granule to those found with RNA Pol II (Thompson et al. 2018; Chong et al. 2018; Wei et al. 2020). We investigated assemblies stabilized by crosslinking for analysis by SEC or IP assays. The LC domain forms protein interactions and is critical to function, including EWS-FLI1 binding to DNA microsatellites (Johnson et al. 2017; Boulay et al. 2017; Kwon et al. 2013). Crosslinking did not alter the ability of mutations to the LC domain to prevent EWS-FLI1 binding to EWSR1 or RNA Pol II (**Figure 5B**). FET protein assemblies vary by appearance *in vitro*, from fibers and aggregates to liquid-like condensates that adopt round shapes or spheres in suspension (Springhower et al. 2020; McSwiggen et al. 2019b; Lin et al. 2015; Molliex et al. 2015). These variant forms may place architectural constraints on interactions with transcription machinery (McSwiggen et al. 2019a; Thompson et al. 2018; Murray et al. 2017; Schwartz et al. 2013). Particles recovered by immunoprecipitation of EWS-FLI1 were observed only as rounded forms (**Figure 5D**). This appearance is consistent with the structures of cellular condensates or granules, rather than fibers or amorphous aggregates (Kwon et al. 2013; Chong et al. 2018). These assemblies may represent at least part of a molecular scaffold that facilitates EWS-FLI1 interactions with the large network partners regulating gene expression in Ewing sarcoma (Gorthi et al. 2018; Boulay et al. 2017; Sankar and Lessnick 2011).

More EWS-FLI1 target genes were affected by the siRNA knockdown of EWSR1 than expected (Sankar et al. 2013). Some functions of EWSR1 are reduced or absent in Ewing sarcoma due to dominant negative effects of EWS-FLI1 (Gorthi et al. 2018; Embree et al. 2009). Since EWS-FLI1 binds EWSR1 through the domains they share and recruits EWSR1 to its binding sites along DNA, it is reasonable to hypothesize that a function remains for EWSR1 that may be incorporated into that of the fusion protein (Boulay et al. 2017; Gorthi et al. 2018). Notable differences in approach may have allowed the new observations made in this study. The siRNA knockdown is relatively brief for the cells, while a stable knockdown by an shRNA reveals effects sustained as cells continue to grow over multiple passages in the absence of EWSR1 (Sankar et al. 2013). The knockdown in HEK293T/17 cells also produced fewer effects on transcript levels. Several observations argue the siRNA knockdown effects were not off target. First, the overlap in changes to transcript levels was not random (**Figure 1**). Both knockdowns affected established EWS-FLI1 targets, both repressed and activated genes (Stoll et al. 2013; Smith et al. 2006; Riggi et al. 2014). Similar GO associations were enriched among genes affected by EWSR1 or EWS-FLI1 in Ewing sarcoma cells, suggesting the overlap in activity has a functional significance (**Supplemental Figure S2**). The results of EWSR1 knockdown by shRNA can teach us that EWSR1 activity is not irreplaceable, and cells can accommodate this loss (Sankar et al. 2013; Kishore et al. 2014). A hypothesis that EWSR1 contributes to heterogeneous protein structures associated with EWS-FLI1 activity may allow that cells adapt through substituting a protein of similar capabilities, which may include FUS, TAF15, or another hnRNP protein. EWSR1 functions that are missing in Ewing sarcoma may offer an easier transition to growth and transcription regulation without this binding partner (Gorthi et al. 2018; Li et al. 2007).

Anchorage-independent growth was also affected by both EWS-FLI1 and EWSR1, but the knockdown in HEK293T/17 suggested EWSR1 may not be universally required for growth of transformed cells (**Figure 2D**). These results add to evidence that EWS-FLI1 can modify EWSR1 activity or vis versa. The acquired sensitivity to the EWSR1 knockdown when exogenous EWS-FLI1 is expressed suggests that the interaction is functionally significant (**Figure 3B**). Recent studies have provided new insights into EWSR1 and EWS-FLI1 interactions that may be particularly relevant. EWS-FLI1 blocks EWSR1 activity that prevents R-loops from either forming or persisting during transcription, which adds to transcriptional and replication stress in Ewing sarcoma (Abraham et al. 2020; Gorthi et al. 2018). We noted that exogenous EWS-FLI1 can raise transcript levels closer to that observed in Ewing sarcoma cells, where transcript levels can be further increased by removal of EWS-FLI1. This may indicate more than one mechanism can drive and moderate expression of EWS-FLI1 targets in Ewing sarcoma. Further investigations of cells that thrive without EWSR1 may reveal which critical functions must be restored for EWS-FLI1 to continue to sustain tumor growth and help identify new factors involved.

A nuclear granule formed by the fusion protein driving Ewing sarcoma raises exciting new questions. How many interactors of EWS-FLI1 possess oligomerization properties like EWSR1? Proteins playing a structural role in EWS-FLI1 assemblies may add new functionality or modify other protein components. Second, are more hnRNP proteins and RNA processing machineries recruited to DNA sites occupied by EWS-FLI1? If true, at least three possibilities can be explored: can RNA Pol II be recruited by the RNA processing machinery itself; would the fusion protein repress transcription by directing RNA processing machinery elsewhere; and does a reorganization of RNA processing machinery contribute to reprogramming transcription toward tumorigenesis. Last, if RNA-binding proteins are recruited to chromatin by the fusion protein, what RNA transcripts can associate with the assemblies and what role might they serve? These questions will likely require knowledge of how many assemblies or granules can incorporate EWS-FLI1, including any that may not be bound to DNA. In future studies, new findings may help rationalize contrasting activities of EWS-FLI1 based on the type of granule or assembly involved.

## MATERIALS AND METHODS

### Cell Culture

Cell lines were obtained from American Type Culture Collection. A673 (ATCC-CRL-1598) and SK-N-MC (ATCC-HTB-10) cells were grown in Dulbecco’s modified Eagle Medium (DMEM, Thermo Fisher) supplemented with 10% fetal bovine serum (FBS). HEK293T/17 (ATCC-CRL-11268) cells were grown in DMEM supplemented with 5% FBS. All cell lines were cultured at 37°C and 5% CO_2_.

### Plasmid and siRNA Transfections

Sequences for siRNAs used are provided in **Supplemental Table 1** and were purchased commercially (Sigma-Aldrich). Oligonucleotides were annealed to form siRNA duplexes by heating to 95 °C in PBS and cooled at room temperature. The plasmids used, pLV-V5-EWS-FLI1 and pLV-V5-EWS(YS37)-FLI1, were a gift from the M. Rivera lab (Harvard Medical School, Charlestown MA). A673 cells (5.0 × 10^5^) in 6-well dishes were reverse transfected with siRNA (50 nM) using Lipofectamine™ RNAiMAX (Invitrogen, cat. no.13778075) according to manufacturer’s instructions. HEK293T/17 and SK-N-MC cells (4.0 × 10^5^) were seeded in 6-well dishes and transfected with siRNA (50 nM) after 24 hours using the TransIT-X2 lipid reagent (Mirus Bio cat. no. MIR6000) or RNAiMAX™. Cells were harvested 48 to 72 hours after siRNA transfections. HEK293T/17 cells grown to 80% confluency for transfection with plasmid (2 μg) using the TransIT-X2 lipid reagent and according to manufacturer’s instructions. Cells transfected with plasmids or plasmid with siRNA were harvested 48 to 96 hours post-transfection.

Knockdown efficiency was assessed for siRNAs screened for use in this study by real time PCR using a StepOne Plus Real-Time PCR System (Applied Biosystems). Total cell RNA was harvested using Trizol™ (Life Technologies) and cDNA synthesized using the High-Capacity cDNA Reverse Transcription Kit (Life Technologies), which uses random primers. Real-time PCR was performed with the TaqMan^®^ Universal PCR Master Mix (Life Technologies). Commercially designed primers (Taqman gene expression assay, Thermo Fisher) were purchased for FLI1 (Hs04334143_m1), EWSR1 (Hs01580532_g1), and the Human GAPD (GAPDH) Endogenous control. Relative change in cDNA was calculated using the ΔΔC_T_ method and normalized to the GAPDH control.

### Western Blot Analysis

Protein lysate concentrations were quantified using the bicinchoninic (BCA) protein assay (ThermoFisher, cat. no. 23227). Protein samples of 5 to 10 μg were loaded onto 7.5% SDS-PAGE gels. Blots were transferred at 500 mA, blocked in 5% nonfat dried milk in Tris-buffered saline-Tween (TBS-T), and incubated overnight with primary antibody at 4°C. Blots were washed in TBS-T, incubated in secondary antibody for 1 hour at room temperature, washed again in TBS-T, and imaged after the addition of SuperSignal™ West Pico PLUS Chemiluminescent substrate (ThermoFisher, cat. no. 34578). Antibodies used in western blots were anti-EWSR1 (clone B-1, Santa Cruz Biotechnology, cat. no. 398318), anti-FLI1 (Abcam, cat. no. 15289), anti-RNA Pol II (clone CTD4H8, EMD Millipore, cat. no. 05-623), anti-RNA Pol II CTD phospho S5 (Abcam, cat. no. 5131), and anti-V5 (Abcam, cat. no. 27671); secondary antibodies used were donkey anti-mouse IgG horseradish peroxidase (HRP) (Jackson ImmunoResearch, cat. no. 715-035-15) and donkey anti-rabbit IgG HRP (Jackson ImmunoResearch, cat. no. 711-035-152).

### RNA Sequencing

A673 or HEK293T/17 cells were transfected with 50 nM siRNA as described above. Cells were collected 72 hours post-transfection and total RNA was extracted using TRIzol reagent (ThermoFisher, cat. no. 15596026). RNA (1 μg) for A673 experiments was used to prepare barcoded sequencing libraries using NEBNext Poly(A) mRNA Magnetic Isolation Module (New England Biolabs, cat. no. E7490) according to the manufacturer’s instructions. A673 libraries (n = 2 per treatment) were sequencing by the University of Arizona Genetics Core (Tucson, AZ) using an Illumina HiSeq2500, yielding between 20M to 32M 100 bp paired end reads. HEK293T/17 libraries were prepared and sequenced using an Illumina HiSeq by Novogene Co., Ltd. (Davis, CA), yielding 17 to 30M 150 bp paired end reads.

FastQ files from this study or publicly available datasets were analyzed essentially as described (Showpnil et al. 2020). Low quality reads and adapters were trimmed using TrimGalore 0.6.6 (https://github.com/FelixKrueger/TrimGalore). Reads were aligned to GRCh38 using STAR 2.7.8a (Dobin et al. 2013). Differential expression analysis used R-studio 4.0.3 libraries available through Bioconductor (Huber et al. 2015). ComBat was used to correct expression for batch effects (Johnson et al. 2007). DESeq2 was used to calculate changes to RNA transcript levels (Love et al. 2014). The top 10706 genes with base mean expression value >60 was classified as “expressed” transcripts. Previously published gene sets for signature repressed or activated EWS-FLI1 targets were analyzed without regard to a threshold for expression (Sankar et al. 2013). Datasets generated by this study are publicly available under GEO Accession no. GSE154944. Alignment statistics for all RNA-seq data analyzed in this study are provide in **Supplemental Table 1**. Fold changes for expressed transcripts and normalized counts are found in **Supplemental Table 2**. Network analysis was performed using Cytoscape v3.7.2 and the BiNGO app using default settings and the ontology files: GO_Cellular_Component, GO_Molecular_Function, and GO_Biological_Process (Excoffier et al. 2017; Shannon et al. 2003). A full list of GO associations found by network analysis is found in **Supplemental Table 3**.

### Soft Agar Colony Formation Assay

A673, SK-N-MC, and HEK293T/17 cells transfected with 50 nM siRNA were harvested at 24 hours posttransfection. HEK293T/17 cells transfected with 50 nM siRNA and 2 μg of plasmid DNA were harvested 24 hours post-transfection. A673 and SK-N-MC cells were seeded at density of 1.0 × 10^5^ cells. Cells were resuspended in 0.35% agarose in growth medium and plated onto a bed of solidified 0.6% agarose in growth media. HEK293T/17 cells were seeded at a density of 20k to 30k cells. Cells were resuspended in 0.4% agarose in growth medium onto a bed of solidified 0.6% agarose in growth media. A673 and SK-N-MC cells were grown at 37°C and 5% CO_2_ for 3 to 4 weeks, imaged, and then colonies were counted using ImageJ software. HEK293T/17 cells were grown at 37°C and 5% CO_2_ for 1 to 2 weeks, imaged, and then colonies were counted. Colonies with stained with 0.05% methylene blue.

### Cell Growth Assay

A673 cells were reverse transfected at a density of 4.0 × 10^5^ cells in 6-well dishes. Cells were collected by trypsinization and counted on a hemocytometer at 24, 48, 72, and 96-hour time points post-transfection.

### Co-Immunoprecipitation Assay

Co-immunoprecipitation assays (co-IP) performed with uncrosslinked cells were grown to confluency in 6-well plates. Cells were harvested and lysed in co-IP lysis/wash buffer (25 mM Tris-HCl pH 7.4, 200 mM NaCl, 1 mM EDTA, 0.5% NP40, 5% glycerol). Protein A/G agarose beads (EMD Millipore, cat. no. IP05) were incubated with primary antibody for 2 hours at 4°C before addition to cell lysate. Lysate was incubated with beads-antibody complex overnight with rotation at 4°C. Beads were washed 5 times in co-IP lysis/wash buffer, resuspended in Novex NuPage™ Sample Loading Buffer (Fisher Scientific, cat. no. np0008) with 5 mM dithiothreitol (DTT) and boiled for 5 minutes at 95 °C. Beads were then centrifuged at 8000 rpm and eluted protein removed with the supernatant and detected by western blot.

For crosslinked co-IP assays, cells were harvested from confluent 150-mm dishes. Cells were crosslinked using 1% formaldehyde for 15 minutes and then quenched with glycine (125 mM). Cells were washed in phosphate-buffered saline (PBS) and lysed in Buffer B (1% SDS, 10 mM EDTA, 50 mM Tris-HCl pH 8.0) supplemented with protease inhibitors. Lysate was sonicated using a Bioruptor Pico (Diagenode) for 30 minutes and then centrifuged at maximum speed (20000×*g*) for 30 minutes at 4°C. Crosslinked lysate was diluted 10-fold in IP lysis buffer (0.01% SDS, 1.1% Triton-X, 1.2 mM EDTA, 16.7 mM Tris-HCl pH 8.0, 167 mM NaCl) treated with protease inhibitors (Sigma-Aldrich, cat. no. P8340) and benzonase (Millipore-Sigma, cat. no. 70746). Lysate was then incubated with rotation overnight with primary antibody at 4°C. Antibody-bound complexes were immunoprecipitated with Novex DYNAL Dynabeads Protein G (Invitrogen, cat. no. 10-003-D) or protein A/G agarose beads (EMD Millipore, cat. no. IP05) for 2 hours at room temperature. Beads were washed 5 times using IP lysis buffer and eluted in 3.6 M MgCl_2_ and 20 mM 2-(*N*-morpholino)ethanesulfonic acid (MES, pH 6.5) for 30 minutes with agitation. IP samples were then assayed for proteins by ELISA.

IP experiments were performed with antibodies to EWSR1 (clone B-1, Santa Cruz Biotechnology, cat. no. 398318), FLI1 (Abcam, cat. no. 15289), RNA Pol II (clone CTD4H8, EMD Millipore, cat. no. 05-623), phospho S5 RNA Pol II (Abcam, cat. no. 5131), and V5 (Abcam, cat. no. 27671). Elutions were probed by western assay for uncrosslinked samples or by ELISA for crosslinked samples.

### Size Exclusion Chromatography

The protocol for SEC of crosslinked lysates has been previously described (Thompson et al. 2018). For crosslinked lysates, cells were harvested from confluent 150-mm dishes. Cells were crosslinked in 1% formaldehyde for 15 minutes and then quenched in 125 mM glycine. Cells were harvested by scraping, washed in PBS, and resuspended in 5 to 10 volumes of Buffer C (400 mM NaCl, 20 mM HEPES pH 7.9, 5% glycerol, 0.75 mM MgCl_2_, and 6 M urea). Lysates were sonicated using a Bioruptor Pico (Diagenode) for 30 minutes at 4°C, followed by centrifugation at maximum speed (20000×*g*) for 30 minutes at 4°C, then filtered through a Costar Spin-X 0.45 μm filter (VWR, cat. no. 8163). Lysate (1 to 2 mg) was injected onto the column. SEC was performed using a Sepharose CL-2B 10/300 column (Sigma-Aldrich, cat. no. CL2B300, 100 mL) injected with lysates from HEK293T/17 or A673 cells. Columns were run in Buffer B (100 mM NaCl, 20 mM HEPES pH 7.9, 0.2 mM EDTA, 5% glycerol, 6 M urea, and 0.5 mM DTT). SEC experiments were analyzed by ELISA as described below with antibodies to EWSR1 (clone C-9, Santa Cruz Biotechnology cat. no. sc-48404) and RNA Pol II (clone CTD4H8, EMD Millipore, cat. no. 05-623).

### ELISA

ELISAs were performed in 96-well Greiner LUMITRAC-600 white plates (VWR, cat. no. 82050-724). ELISAs were performed as indirect ELISAs, in which protein samples from SEC or crosslinked IP assays were incubated in plates overnight at 4°C. Afterward, plates were washed 3 times in TBS-T, blocked for 2 hours at room temperature in 5% nonfat dried milk in TBS-T, washed 4 times in TBS-T and then incubated with primary antibody overnight at 4°C. After incubation with primary antibody, plates were washed 4 times in TBS-T and incubated with secondary antibody, goat anti-mouse IgG HRP (ThermoFisher cat. no. 31432) or goat anti-rabbit IgG HRP (ThermoFisher, Cat. #31462) for 1 hour at room temperature. Finally, plates were washed 4 times with TBS-T and proteins were detected by addition of SuperSignal™ ELISA Femto substrate (ThermoFisher cat. no. PI37074). Luminescence was read using a BMG POLARstar Omega plate reader or Biotek Neo2 microplate reader.

### Transmission Electron Microscopy

TEM assays were performed for samples from co-IP assays using antibodies to RNA Pol II (CTD4H8) or FLI1 (Abcam, cat. no. 15289). Carbon Film 300 Mesh Copper grids (Electron Microscopy Sciences, cat. no. CF300-CU) were charged at 15 mA for 30 seconds. Crosslinked immunoprecipitation samples were spotted onto charged grids and stained with 0.75% uranyl formate. For proteinase K-treated samples, the samples were treated with 5 μg of proteinase K and incubated for 30 minutes at 37°C before being spotted onto grids and stained with 0.75% uranyl formate. Images were collected from a FEI Tecnai Spirit 120S or FEI Tecnai F20 transmission electron microscope. Diameters of particles observed were measured with ImageJ software (U. S. National Institutes of Health, Bethesda, Maryland, USA, https://imagej.nih.gov/ij/).

## Corresponding Author

Correspondence should be addressed to Jacob C. Schwartz at.

## Author Contributions

N.S.A. designed and performed experiments, analyzed data, and interpreted results. L.M.H. contributed the design of siRNAs, performed the screening and optimization siRNA knockdown using real-time PCR and western assays, and contributed to multiple procedures involving recombinant DNA. M.A.L. and V.F.T. prepared samples for RNA-seq analysis and D.R.W. performed analysis and quantification of sequencing data. N.S.A. wrote the first draft of the manuscript. J.C.S. contributed to analysis of RNA-seq data. N.S.A. and J.C.S. collaborated to write subsequent drafts. All authors have approved to the final version of the manuscript.

## Funding Sources and Acknowledgements

This work was supported by funding from the National Institutes of Health (R21CA238499) and the American Cancer Society (RSG-18-237-01-DMC) to J.C.S. Additional support was provided by National Institute of General Medicine (T32-GM008659) training grant awarded to N.S.A. Next-generation sequencing was performed by the University of Arizona Genetics Core, http://uagc.arizona.edu. Transmission election microscopy was performed with the Eyring Materials Center at Arizona State University, supported in part by NNCI-ECCS-1542160. Transmission Electron Microscopy was also performed through the University of Arizona Office of Research, Innovation, & Impact Imaging Cores. Research reported in this publication was also supported by the Office of the Director, National Institutes of Health of the National Institutes of Health, under award number S10OD013237.

