## Supplemental Figures for "Fusion protein EWS-FLI1 is incorporated into a protein granule in cells"

### Supplemental Figure S1

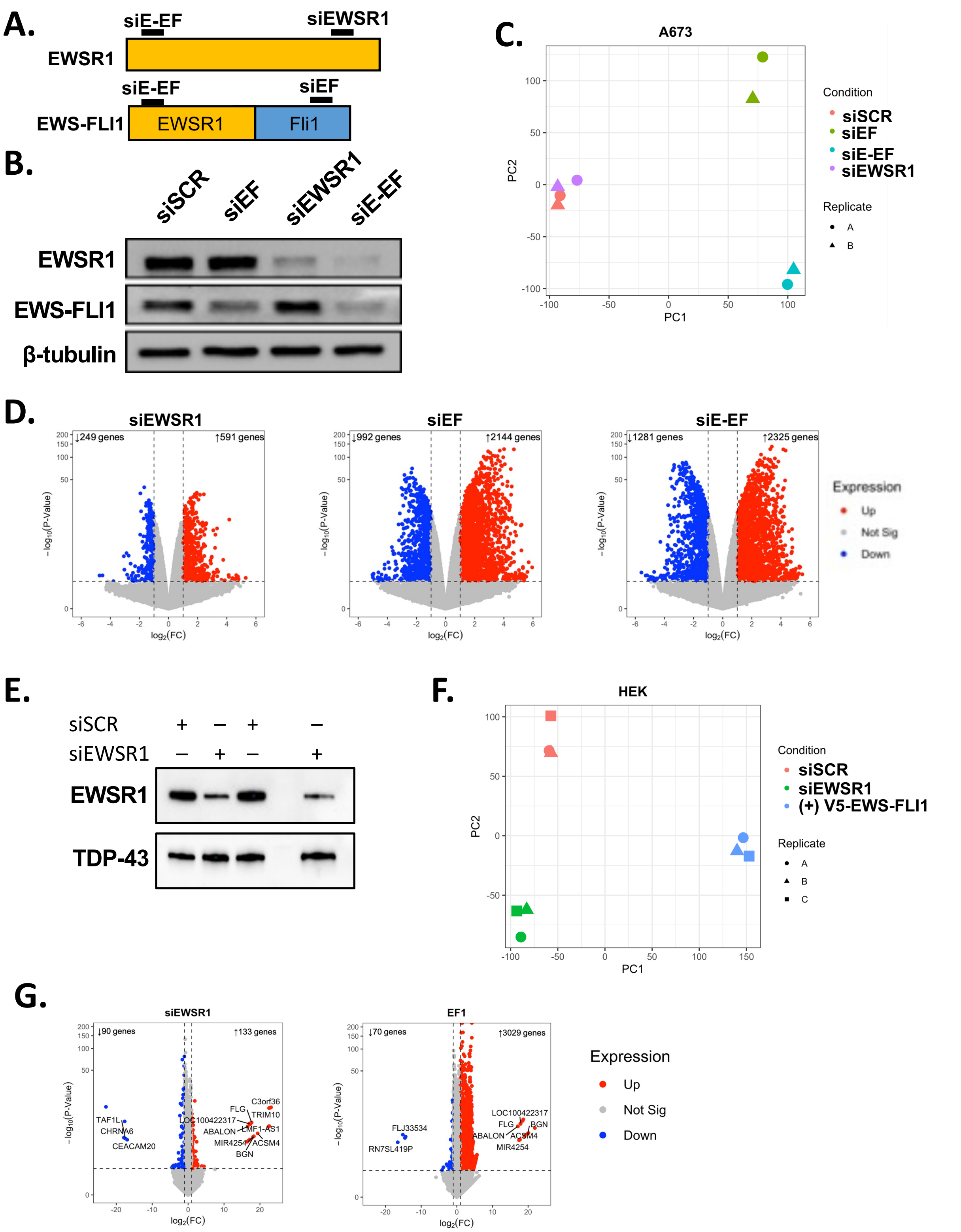

### Supplemental Figure S1 *(continued)*

**H.**

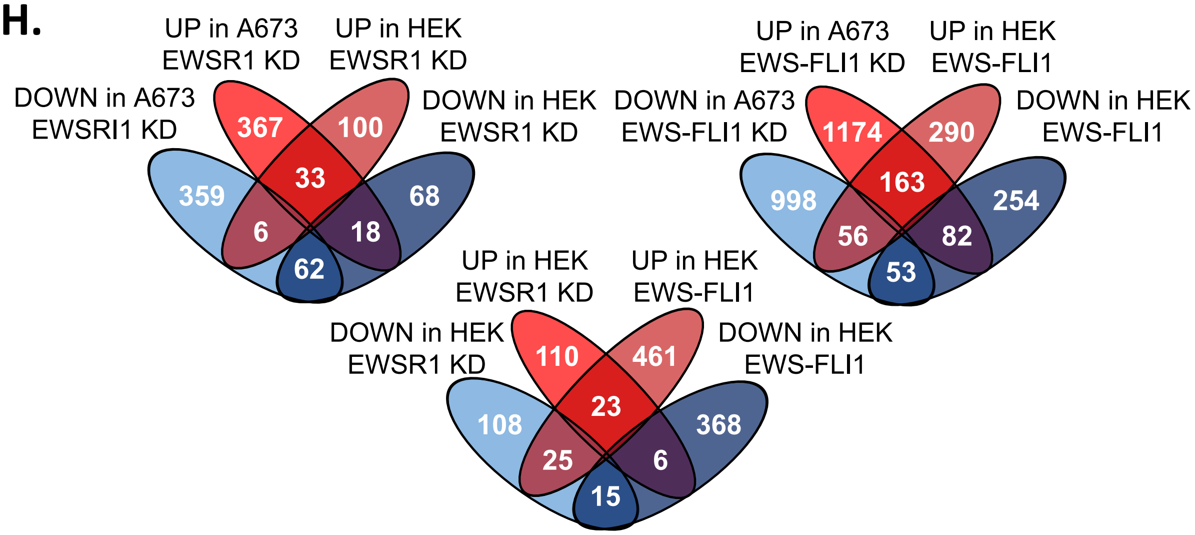


**Figure S1. (A)** Targets for three siRNAs knocking down EWSR1, EWS-FLI1, or both are shown. (B) Western analysis of knockdown in A673 cells for EWSR1 or EWS-FLI1 by siRNA with β-tubulin as a loading control. (C) Principal component analysis of knockdown in A673 of EWSR1, EWS-FLI1, or both, compared to the siSCR control treatment. (D) DESeq2 volcano plots indicating A673 transcripts increased or decreased >2-fold, p-adj <0.05, and no minimum threshold for expression. (E) Western analysis of EWSR1 knockdown in HEK293T/17 with TDP-43 shown as a loading control. (F) Principal component analysis of RNA-seq results for EWSR1 knockdown or exogenous EWS-FLI1 expression in HEK293T/17. (G) DESeq2 volcano plots indicating HEK293T/17 transcripts increased or decreased >2-fold, p-adj <0.05, and no minimum threshold for expression. (H) Vinn diagrams showing the overlap of genes activated or silenced in response to treatment.

### Supplemental Figure 2

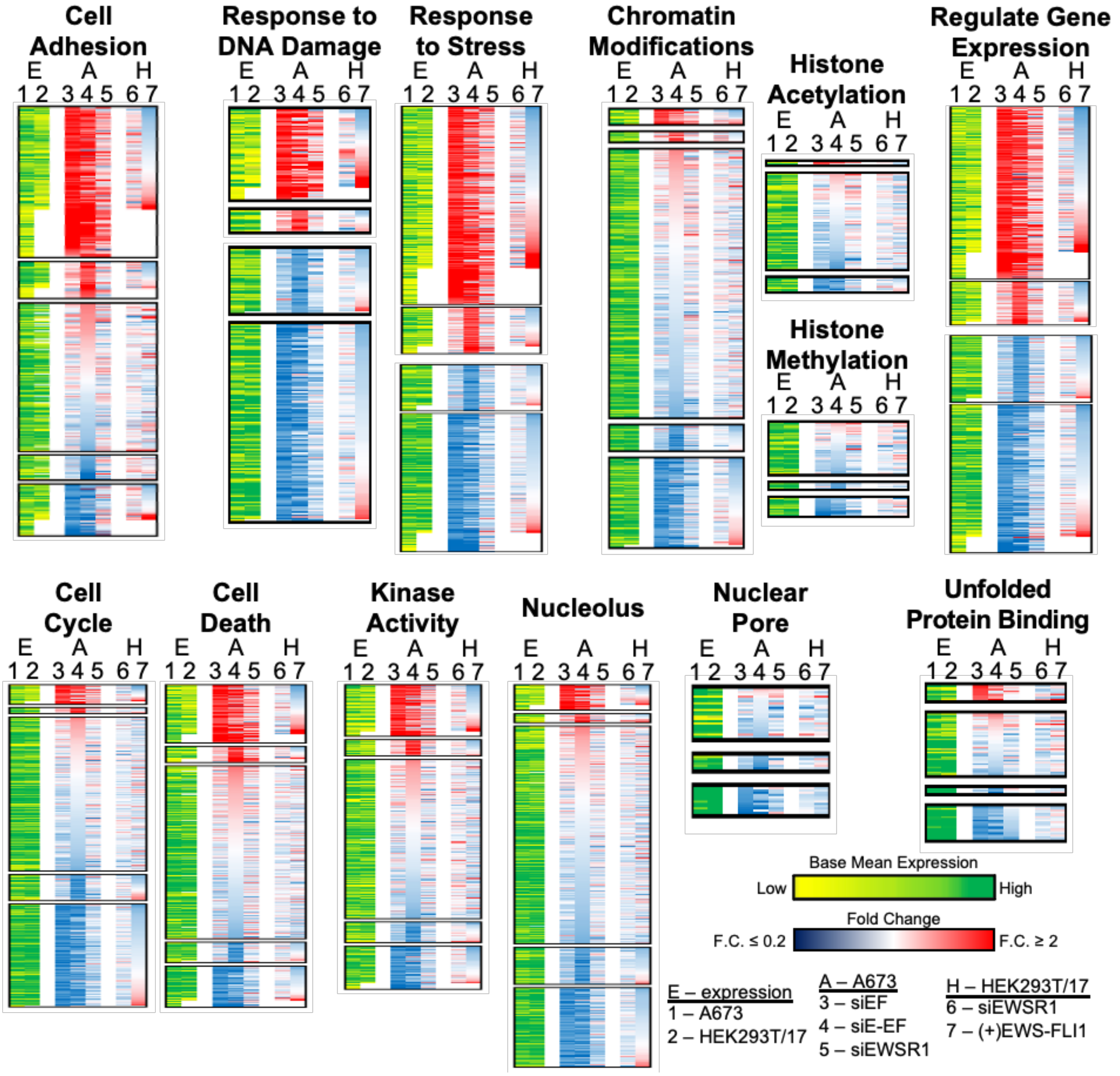


| | | | **siEF** (A673) | | | **siEF & siEEF** (A673) | | | **siEWSR1** (A673) | | | **siEWSR1** (HEK293T/17) | | | **(+)EWS-FLI1** (HEK293T/17) | | |
|---|---|---|---|---|---|---|---|---|---|---|---|---|---|---|---|---|---|
| **UP** | **GO ID** | **Description** (N expressed in A673) | **N (↑)** | **N (↓↑)** | **p-adj** | **N (↑)** | **N (↓↑)** | **p-adj** | **N (↑)** | **N (↓↑)** | **p-adj** | **N (↑)** | **N (↓↑)** | **p-adj** | **N (↑)** | **N (↓↑)** | **p-adj** |
| | 3700 | **transcription factor activity** (585) | 112 | 212 | 4E-03 | 143 | 290 | 6E-03 | 60 | 81 | 1E-02 | **12 (↓)** | 15 | NA | 41 | 81 | 1E-07 |
| | 7155 | **cell adhesion** (510) | 184 | 248 | 1E-22 | 232 | 328 | 1E-13 | 77 | 113 | 5E-03 | 12 | 20 | NA | 28 | 38 | NA |
| | 6950 | **response to stress** (1841) | 374 | 656 | 1E-11 | 463 | 841 | 1E-09 | 178 | 272 | 1E-06 | 26 | 52 | NA | 87 | 110 | 2E-02 |
| | 10468 | **regulation of expression** (2643) | 380 | 756 | 8E-03 | 477 | 1042 | 6E-06 | 181 | 307 | NA | 22 | 49 | NA | 111 | 168 | 1E-04 |
| | 8219 | **cell death** (549) | 102 | 175 | 2E-02 | 130 | 239 | 5E-08 | 58 | 84 | 2E-04 | **9 (↓)** | 11 | NA | 23 | 31 | NA |
| | 16301 | **kinase activity** (524) | 93 | 171 | NA | 121 | 237 | 4E-04 | 31 | 67 | NA | 8 | 11 | NA | 23 | 30 | NA |
| **DOWN** | **GO ID** | **Description** (N expressed in A673) | **N (↓)** | **N (↓↑)** | **p-adj** | **N (↓)** | **N (↓↑)** | **p-adj** | **N (↓)** | **N (↓↑)** | **p-adj** | **N (↓)** | **N (↓↑)** | **p-adj** | **N (↓)** | **N (↓↑)** | **p-adj** |
| | 6281 | **DNA repair** (385) | 95 | 113 | 1E-16 | 124 | 147 | 4E-27 | 15 | 20 | NA | 1 | 3 | NA | **17 (↑)** | 23 | NA |
| | 6974 | **response to DNA damage** (594) | 125 | 181 | 1E-19 | 167 | 238 | 5E-29 | 26 | 51 | NA | 6 | 9 | NA | **27 (↑)** | 32 | NA |
| | 16568 | **chromatin modification** (403) | 84 | 101 | 1E-03 | 113 | 139 | 1E-08 | 20 | 32 | NA | 2 | 3 | NA | **8 (↑)** | 11 | NA |
| | 16571 | **histone acetylation** (89) | 10 | 13 | NA | 18 | 22 | NA | 4 | 6 | NA | NA | 0 | NA | **1 (↑)** | 1 | NA |
| | 16573 | **histone methylation** (61) | 14 | 14 | NA | 19 | 19 | NA | 5 | 7 | NA | 1 | 1 | NA | 2 | 2 | NA |
| | 7049 | **Cell cycle** (548) | 180 | 215 | 1E-42 | 227 | 273 | 2E-56 | 31 | 43 | NA | 3 | 3 | NA | **9 (↑)** | 15 | NA |
| | 5643 | **nuclear pore** (46) | 14 | 15 | 2E-03 | 21 | 22 | 2E-07 | 6 | 6 | NA | 1 | 1 | NA | **2 (↑)** | 2 | NA |
| | 5730 | **nucleolus** (654) | 137 | 183 | 5E-06 | 204 | 265 | 2E-29 | 36 | 55 | NA | 5 | 7 | NA | **19 (↑)** | 27 | NA |
| | 51082 | **unfolded protein binding** (77) | 21 | 32 | NA | 26 | 38 | 2E-03 | 6 | 9 | NA | **1 (↑)** | 1 | NA | **3 (↑)** | 3 | NA |

**Figure S2.** Heatmaps are shown for GO associated genes listed in the table. Heat maps are sorted by changes in transcripts abundance after knockdown in A673 by siEF (3), then siE-EF (5), and finally exogenous expression of EWS-FLI1 in HEK293T/17 (7). Also shown are base mean expression values for genes in A673 (1) or HEK293T/17 (2).

### Supplemental Figure 3

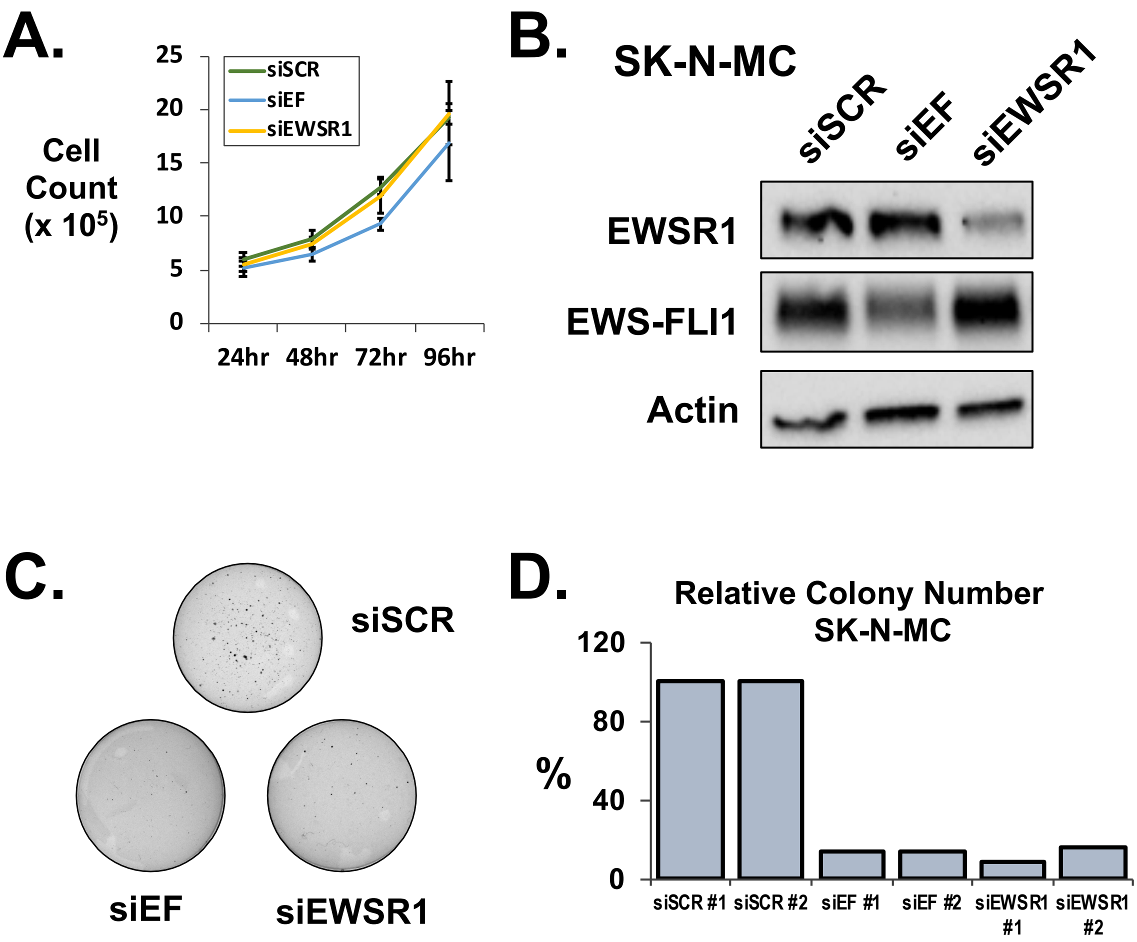


**Figure S3. A.** Cell growth assay of A673 after transfection of siRNAs targeted against EWS-FLI1 (siEF), EWSR1 (siEWSR1), or a scramble control (siSCR). **B.** Western blot of knockdown of EWS-FLI1, EWSR1, and scramble control in SK-N-MC cells. **C.** Soft agar assays in SK-N-MC cells following knockdown of EWS-FLI1, EWSR1, or scramble control. **D.** Bar graph quantifies relative colony count in SK-N-MC cells. (n=2).

### Supplemental Figure 4

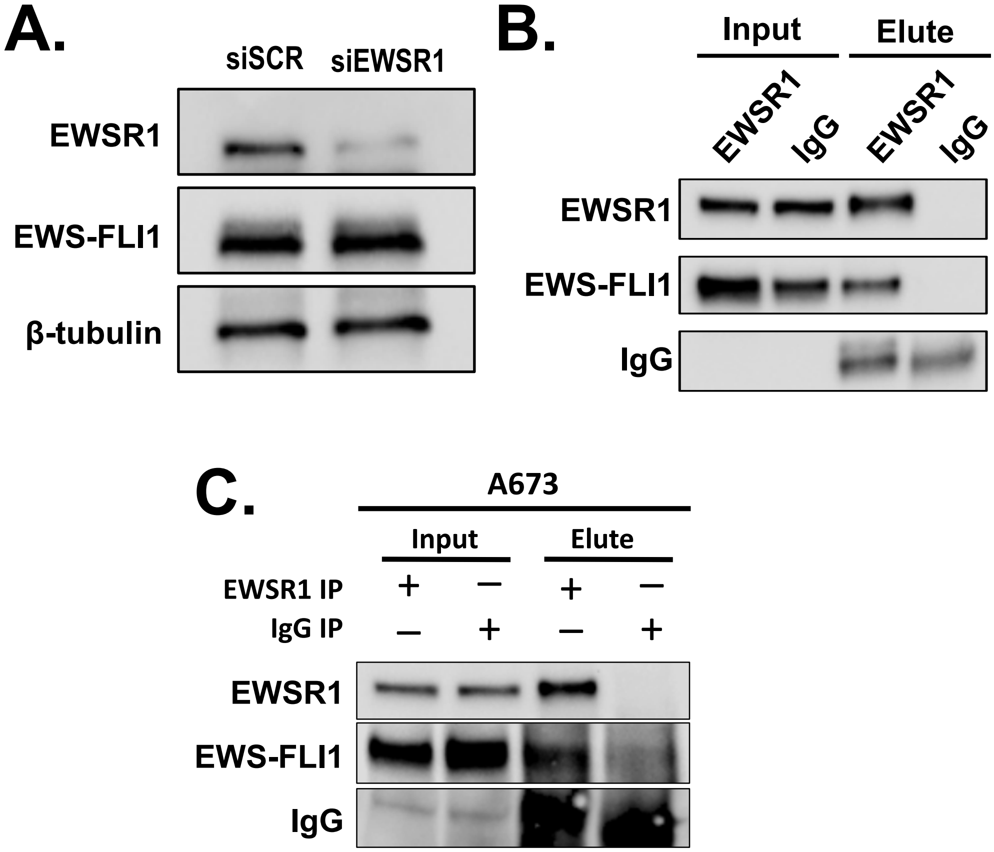


**Figure S4. A.** Western blot of HEK293T/17 cells co-transfected with V5-tagged EWS-FLI1 (V5 EWS-FLI1) and siRNA targeting EWSR1 or scramble control. **B.** Western blot of co-immunoprecipitation of EWS-FLI1 with EWSR1 (B-1) from HEK293T/17 cells transfected with V5-EWS-FLI1. V5-EWS-FLI1 was detected using anti-V5 antibody. Mouse IgG serves as the negative control. **C.** An immunoprecipitation (n = 1) of endogenous EWS-FLI1 with EWSR1 (B-1) from A673 cells. For this assay, 3 additional replicates did not yield sufficient signals in the EWSR1 IP to reach significance above background signals for the IgG control.

### Supplemental Figure 5

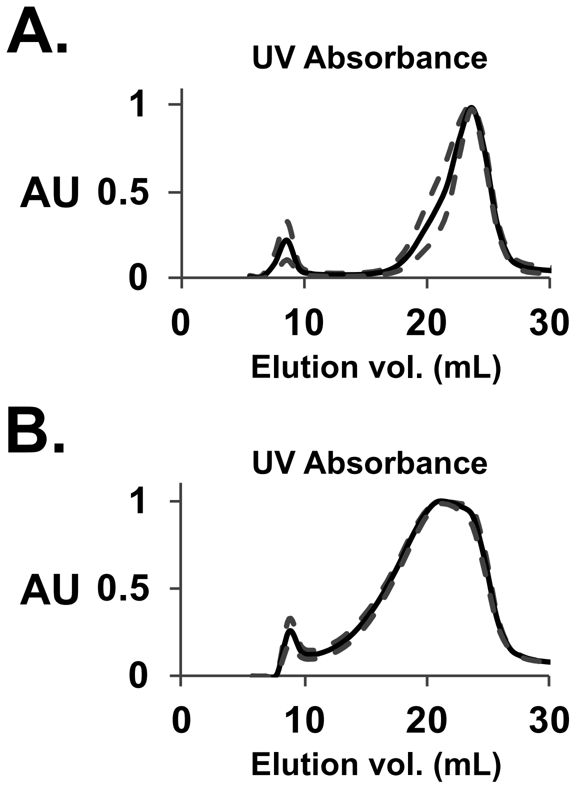


**Figure S5. A.** UV absorbance of total protein of uncrosslinked HEK293T/17 lysates (n=3). **B.** UV absorbance of total protein of crosslinked HEK293T/17 lysates (n=3). Dashed lines represent standard error about the mean.

### Supplemental Figure 6

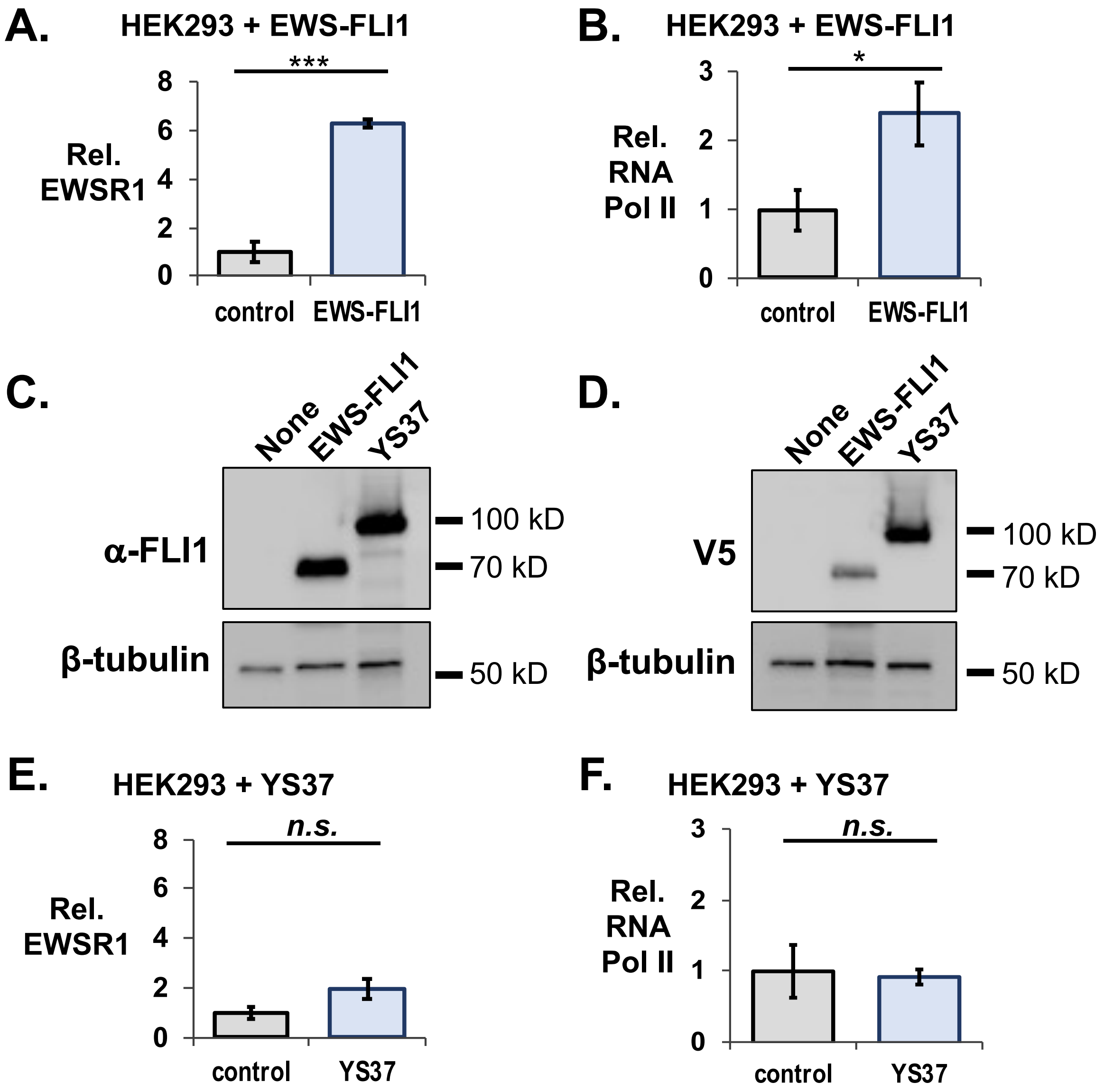


**Figure S6. A.** Crosslinked immunoprecipitation assay of EWS-FLI1 from HEK293T/17 cells transfected with EWS-FLI1 as assessed by ELISA. Endogenous EWSR1 immunoprecipitates with EWS-FLI1. **B.** Crosslinked immunoprecipitation assay of EWS-FLI1 from HEK293T/17 cells transfected with EWS-FLI1 as assessed by ELISA. Endogenous RNA Pol II immunoprecipitates with EWS-FLI1. Western blot of HEK293T/17 cells with no transfection control, EWS-FLI1, and EWS(YS37)-FLI1 probed with **C.** an antibody targeted against the C-terminal FLI1 domain and **D.** an antibody targeted the V5-tag on the N-terminal domain of EWS-FLI1. **E.** Crosslinked immunoprecipitation assay of EWS-FLI1 from HEK293 cells transfected with YS37 EWS-FLI1 mutant as assessed by ELISA. There is no significant enrichment of endogenous EWSR1 with the YS37 mutant. **F.** Crosslinked immunoprecipitation assay of EWS-FLI1 from HEK293 cells transfected with YS37 EWS-FLI1 mutant as assessed by ELISA. There is no significant enrichment of endogenous RNA Pol II with the YS37 mutant. Error bars represent standard error. Student's T-test, *** = $p<0.005$; * = $p<0.05$; n.s. = $p>0.05$.

### Supplemental Figure 7

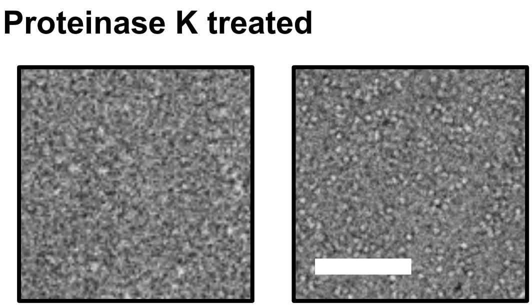


**Figure S7.** Transmission electron microscopy images of crosslinked immunoprecipitated RNA Pol II treated with 5 ug of proteinase K. Scale bar = 50 nm.
